# Proteomic comparison of epidemic Australian *Bordetella pertussis* biofilm cells

**DOI:** 10.1101/2024.02.19.581093

**Authors:** Hiroki Suyama, Laurence Don Wai Luu, Ling Zhong, Mark J. Raftery, Ruiting Lan

## Abstract

*Bordetella pertussis* causes whooping cough, a severe respiratory infectious disease. Studies have compared the currently dominant single nucleotide polymorphism (SNP) cluster I (pertussis toxin promoter allele, *ptxP3*) and previously dominant SNP cluster II (*ptxP1*) strains as planktonic cells. Since biofilm formation is linked with *B. pertussis* pathogenesis *in vivo*, this study compared the biofilm formation capabilities of representative strains of cluster I and cluster II. Confocal laser scanning microscopy found that the cluster I strain had a denser biofilm structure compared to the cluster II strain. Differences in protein expression of the biofilm cells were then compared using Tandem Mass Tagging (TMT) and high-resolution multiple reaction monitoring (MRM-hr). In total, 1453 proteins were identified of which 40 proteins had significant differential expression between the two strains in biofilm conditions. Of particular interest was a large increase in expression of energy metabolism proteins (cytochrome proteins PetABC and BP3650) in the cluster I strain. When the expression of these proteins was compared between 6 additional strains from each cluster, it was found that the protein expression varied between all strains. These findings suggest that there are large levels of individual proteomic diversity between *B. pertussis* strains in biofilm conditions despite the highly conserved genome of the species. Overall, this study revealed visual differences in biofilm structure between *B. pertussis* strains and highlighted strain specific variation in protein expression that dominate potential cluster specific changes that may be linked with the dominance of cluster I strains.

**Importance:** *Bordetella pertussis* causes whooping cough. The currently circulating cluster I strains have taken over previously dominant cluster II strains. It is important to understand the reasons behind the evolution to develop new strategies against the pathogen. Recent studies have shown that *B. pertussis* can form biofilms during infection. This study compared the biofilm formation capabilities of a cluster I and a cluster II strain and identified visual differences in the biofilms. The protein expression between these strains grown in biofilms were compared and proteins identified with varied expression were measured with additional strains from each cluster. It was found that despite the highly conserved genetics of the species, there was varied protein expression between the additional strains. This study highlights that strain specific variation in protein expression during biofilm conditions that may dominate the cluster specific changes that may be linked to the dominance of cluster I strains.

## Introduction

Whooping cough (pertussis) is a severe respiratory infection caused by the bacterium, *Bordetella pertussis.* Despite widespread global vaccination, a resurgence in pertussis cases has been observed globally (1–3). A contributing factor for the resurgence is the evolution of strains with greater fitness and mismatches with the vaccine allele types (3, 4). In Australia, a whole cell vaccine (WCV) was introduced in the 1950s but due to its reactogenicity was subsequently replaced by the acellular vaccine (ACV) in 2000 (5). *B. pertussis* is a homogeneous species with a low level of genetic diversity in terms of single nucleotide polymorphisms (SNPs) (6–8). Our studies have shown that strains belonging to SNP cluster I containing the pertussis toxin promoter *ptxP3* allele has overtaken the previously dominant SNP cluster II strains containing the pertussis toxin promoter *ptxP1* allele (6). Interestingly, studies using a mouse model showed that the cluster I strains had greater fitness compared to the cluster II strains regardless of the immunisation status of the host signifying that there are potentially further changes other than antigenic mismatches with the ACV that allows for the increased fitness of the cluster I strains (9).

Transcriptomic studies have shown that there were a large number of genes with altered expression between *ptxP3* and *ptxP1* strains (4). Major changes in virulence factors, including an increase in pertussis toxin production, were identified and hypothesised to increase colonisation in mice (4, 10). Furthermore, proteomic studies have identified other changes that have occurred between SNP cluster I and SNP cluster II strains (11–13). Changes in expression of proteins associated with immune evasion, virulence and metabolism have been identified in planktonic cells and their secretome (12). Of key importance, it was seen that there was a decrease in the secretion of type 3 secretion system (T3SS) proteins in cluster I strains which may be related to a change from T3SS allele *bscI1* to *bscI3* (11, 12, 14). These additional changes may be associated with the increased fitness observed in the cluster I strains.

Recent studies have shown that biofilms play a key role in *B. pertussis* pathogenesis (15, 16). Biofilms are a community of cells that are encased within a matrix and cells within biofilms have been shown to be more resilient against antimicrobials and environmental stress (17–19). *B. pertussis* has been shown to form biofilms *in vivo* in the upper respiratory tract of a mouse model and strains that lack the ability to form biofilms were less effective at colonisation (15, 20, 21). Furthermore, increased biofilm formation capability has been linked to increased colonisation ability (20). Clinical *B. pertussis* isolates have been shown to form increased biofilm compared to the Tohama I reference strain isolated in the 1950s (17, 20, 22). These studies have demonstrated that biofilms are an important aspect of *B. pertussis* pathogenesis.

Previous studies have identified expression differences between cluster I *ptxP3* and cluster II *ptxp1* strains under planktonic conditions (4, 10–12, 23, 24). However, cells in biofilms are known to be phenotypically distinct from their planktonic counterpart with major changes in transcriptomic and proteomic profiles (17, 22, 25–30). Therefore, comparison between the cluster I and cluster II strains in biofilm conditions may provide additional insights into the adaptation of cluster I strains and re-emergence of *B. pertussis*. This study examined biofilm forming capabilities and proteomic changes between cluster I and cluster II biofilms to identify variations that may help explain the increased fitness of the cluster I strains.

## Methods

### Strains and growth conditions

Strains used in this study are listed in Tables. Clinical isolates, L1423 and L1191, were used as representatives of SNP cluster I and SNP cluster II, respectively. These strains have been previously sequenced and used for comparative infection and proteomic studies (9, 12). The additional strains used to confirm changes in the clusters are all from different years but classified as the same SNP profile (SP) as defined by Octavia *et al.* (6). All cluster I strains are SP13 spanning from the 2008 – 2010 outbreak in Australia while all strains in cluster II are SP39 ranging from 2000 – 2006 (Tables). *B. pertussis* strains were grown on Bordet-Gengou (BG, BD Scientific) agar for 3-5 days at 37°C. For liquid culture, a loopful of pure Bvg^+^ colonies were suspended in 20 mL of THIJS media (31) supplemented with heptakis ((2,6-O-dimethyl) β-cyclodextrin) and 1% THIJS supplement and grown for 24 h shaking at 180 rpm at 37°C.

### Confocal microscopy

Confocal microscopy was used to identify structural changes that occur over time and to identify biofilm maturation. Microscopy was performed on the two representative strains from cluster I (L1423) and II (L1191). A previously established protocol was used with minor changes (32). Briefly, the liquid culture described above was adjusted to OD_600_ of 0.1/mL and seeded into 24 well polystyrene tissue culture plates (Corning) with 12 mm circular sterile glass coverslips (Livingstone, Thickness: No. 1) on the base. The plate was incubated at a 45° angle so the liquid to air interface was localised on the coverslip. The plate was incubated statically for 5 h to allow attachment of cells before the media was refreshed (25, 29). The plates were further incubated at 37°C for 24, 48, 72 and 96 h with mild agitation at 60 rpm (33). At each time point, the wells were washed 3 times with 1 X PBS and the coverslip carefully removed from the well. The biofilm was fixed with 4% paraformaldehyde and stained with SYTO 9 fluorescent dye (Thermo Fisher Scientific). Z-stack images were captured on the Olympus FluoView FV1200 confocal microscope. Three fields of view were randomly selected per sample. Each field of view covering a surface area of 84.48 x 84.48 µm was imaged with Z-stacks (0.42 µm/slice). All images were imaged at 60X objective (oil immersion, NA 1.35; Olympus). The COMSTAT2 (v 2.1) plugin on ImageJ (v 2.8.0) was used to calculate biomass, surface area and maximum thickness of biofilms (34). Three biological replicates per strain per time point were performed.

### Biofilm protein extraction

A previously established method for protein extraction from *B. pertussis* cells (12, 30) was used with minor changes. Briefly, all wells of a 24 well plate were seeded with a 24 hr liquid culture as described above. Each plate represented a single biological replicate, and 7 biological replicates were performed per strain. The plates were incubated statically for 5 hr at 37°C to allow cell attachment before the media was refreshed. The plates were then incubated for a further 96 h shaking at 60 rpm. At 96 h, the supernatant was removed, and the wells washed 3 times with sterile 1 X PBS. Disruption buffer (50 mM Tris-HCl, 0.4 mM phenylmethylsulfonyl fluoride (PMSF) and 2mM EDTA) was added to inactivate proteases. The plates were water bath ultrasonicated at 37 kHz for 2 mins to detach the biofilm cells. All the wells from each individual plate were pooled, centrifuged and resuspended in disruption buffer. The samples were then probe sonicated and proteins extracted using a method previously described by Luu *et al.* (12, 30). A Qubit Fluorometer (Thermo Fisher Scientific) was used to quantify the protein concentration.

### Protein digestion and TMT labelling

For Tandem Mass Tagging (TMT), 100 µg of protein from each replicate was reduced with dithiothreitol, alkylated with iodoacetamide and trypsin digested as previously described by Luu *et al.* (35). Samples were cleaned up using Empore high performance extraction disk cartridges (3M). Following the clean-up, the samples were labelled using the TMT10plex and TMT6plex labels as described in the manufacturer’s protocol (Thermo Fisher Scientific). To account for 7 biological replicates per strain, a 10plex and 6plex kit was used separately. A pooled reference with all samples was included as a reference for normalisation in both experiments. After labelling the peptides with TMT, a styrene divinylbenzene (SDB) stage tip was used to clean up excess tags. The samples were dried using a Savant SpeedVac (Thermo Fisher Scientific). Samples were resuspended in 0.1% formic acid then loaded onto the QExactive Orbitrap mass spectrometer coupled with an UltiMate 3000 high performance nano liquid chromatography system (Thermo Fisher Scientific). Peptides were eluted over a 240 min gradient, ramping from H_2_O:CH_3_CN (98:2, 0.1% formic acid) to H_2_O:CH_3_CN (64:36, 0.1% formic acid) and introduced directly with electrospray ionisation with a flow rate of 200 nL/min. MS spectra were obtained across the mass range of 350 – 1,750 m/z at a resolution of 70,000 m/z, and MS/MS resolution at 45,000 m/z. The 15 most abundant ions were selected for fragmentation with higher energy collision disassociation (HCD). Two technical replicates were performed for each biological replicate.

### Peptide identification and quantification

The TMT10 and TMT6 experiments were combined and normalised to the pooled reference sample. The MS spectra was imported into the Proteome Discover (Thermo Fisher Scientific) software (v2.3) and searched using SEQUEST HT against a custom *B. pertussis* database containing complete genomes of Tohama I, CS, B1917 (*ptxP3*) and B1920 (*ptxP1*). Peptide tolerance was set at 4 ppm and the MS/MS tolerance at 0.04 Da. Variable modifications were set as: Carbamidomethyl (C) and Oxidation (M) and fixed modifications as: TMT-6plex (K), TMT-6plex (N-term), TMT-10plex (K) and TMT-10plex (N-term). Enzyme specificity was set as trypsin with a maximum of 1 missed cleavage and minimum peptides per protein set as 2. Proteins with fold change (FC) > 1.2 and *p* < 0.05 (Benjamini-Hochberg (BH) corrected) were considered upregulated and FC < 0.8 and *p* < 0.05 (BH) were considered downregulated. These cut-offs are widely used by other proteomic studies (35). Functional categories were assigned based on Bart *et al*. (2). pSORTb (v3.0.2) was used to predict subcellular locations of each protein.

### MRM-hr

To confirm the changes in protein expression identified in the TMT-MS, high resolution multiple reaction monitoring (MRM-hr) was used. MRM-hr experiments were run on the two representative strains (L1423 and L1191) and then further performed on 6 additional strains from each cluster. Control proteins were selected as internal standards to normalise the intensities. These proteins were selected based on an FC = 1 ± 0.1 in the TMT experiment. The analysis was performed on the QExactive Orbitrap mass spectrometer. The analysis was designed using Skyline (v21.1.0.278). Protein peptides were screened for suitability for MRM-hr. Peptides with potential ragged ends, missed cleavages or amino acids susceptible to variable modifications were excluded from the analysis. Proteins were excluded unless there was a minimum of 2 suitable peptides identified. The highest peak height and area were selected for each peptide based on the Skyline automated peak detection method (36). Six biological replicates were performed for the representative strains while each additional cluster strain was considered a biological replicate for the cluster. Two technical replicates per biological replicate was injected into the mass spectrometer for analysis.

Ten micrograms of protein were trypsin digested as described above. Samples were cleaned up using C18 StageTips (Thermo Fisher Scientific) and injected into the mass spectrometer over a 30 min gradient. The MRM-hr acquisition consisted of a 50 ms MS scan followed by MS/MS scans of 35 candidate ions per cycle (110 ms accumulation time, 35,000 m/z resolution). The collision energy for each peptide was determined using Skyline and is available in Supplementary Table S1. Retention times were scheduled based on pilot non-scheduled MRM-hr runs. The results were analysed on Skyline using MS-stats (v4.1.2), an inbuilt R package (37).

### Genome comparison of L1423 and L1191

Complete genomes of *B. pertussis* strains, L1423 (SNP cluster I) and L1191 (SNP cluster II) have previously been sequenced (9). The genome sequences were compared using Mauve (v20150226) (38). Protein sequences were compared using Clustal Omega (39).

## Results

### Comparison of biofilm formation between SNP cluster I and SNP cluster II strains

Confocal microscopy was performed to identify changes in biofilm formation between a cluster I strain, L1423 and a cluster II strain, L1191. We confirmed that *B. pertussis* formed mature biofilms at 96 h as previously reported (15, 20, 30). While the biofilm formation over time was similar between the two strains, there were clear visual differences observed between the L1423 and L1191 strains at 96 h (Figure 1). The L1423 strain showed a denser structure compared to L1191. Using the COMSTAT2 plugin on ImageJ, biomass, average thickness and maximum thickness were measured for each strain. Despite the visual differences in structure at 96 h, there was no significant differences in biofilm average thickness, maximum thickness, biomass or roughness coefficient between the two strains at any time point (Figure 2).

**Figure 1.**
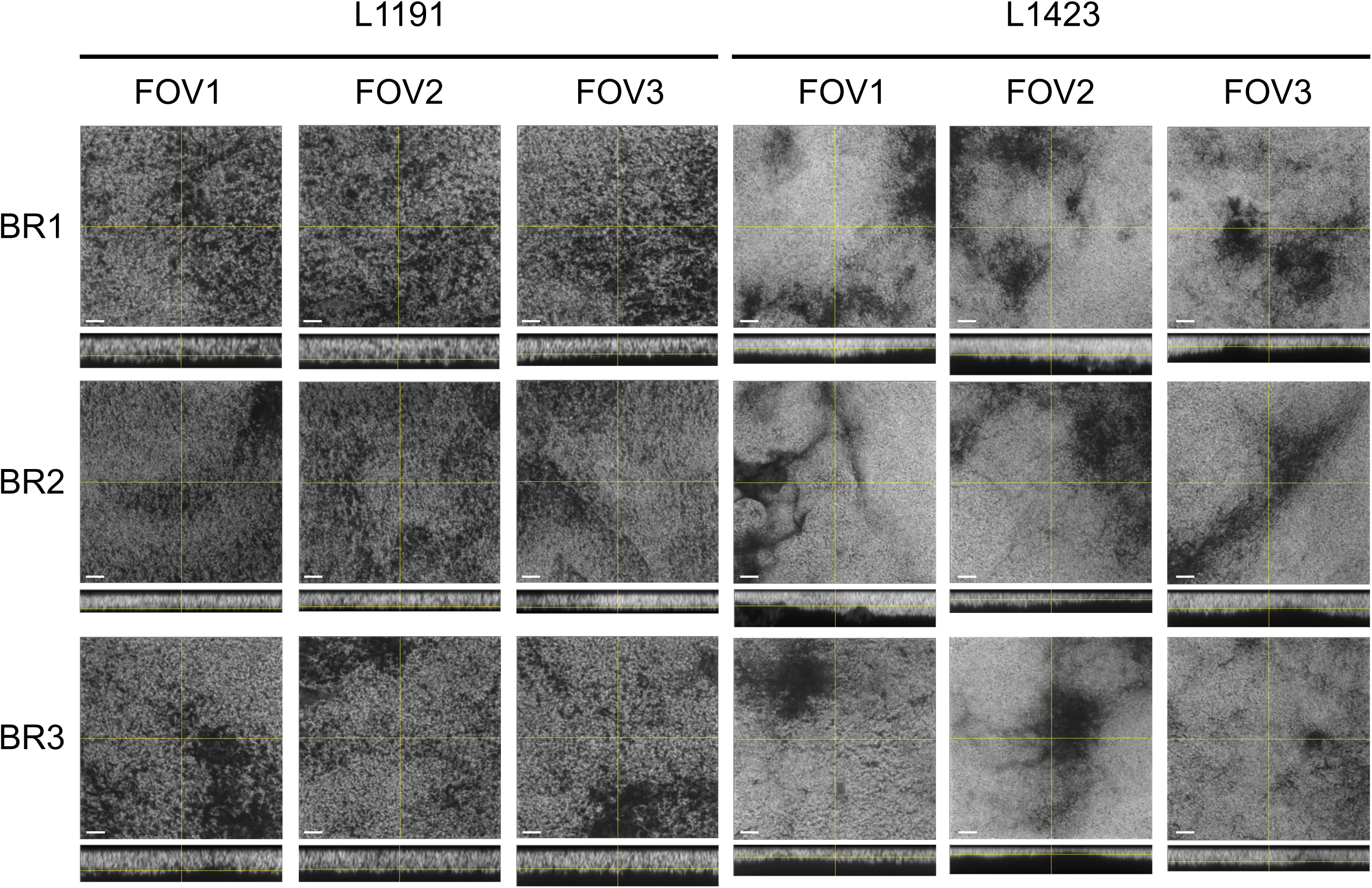
CLSM micrographs of *B. pertussis* biofilms at 96 h. SNP cluster I strain, L1423, and SNP cluster II strain, L1191, were grown on glass cover slips and imaged at 96 h. Biofilms were stained with SYTO9 fluorescent dye and visualised using CLSM. Z-stack images were taken at 0.42 µm intervals. The bar represents 10 µm distance; *xy* and *xz* representative focal planes are shown. Images are represented in original grayscale for easier visibility of contrast (60). BR = biological replicate; FOV = Field of view.

**Figure 2.**
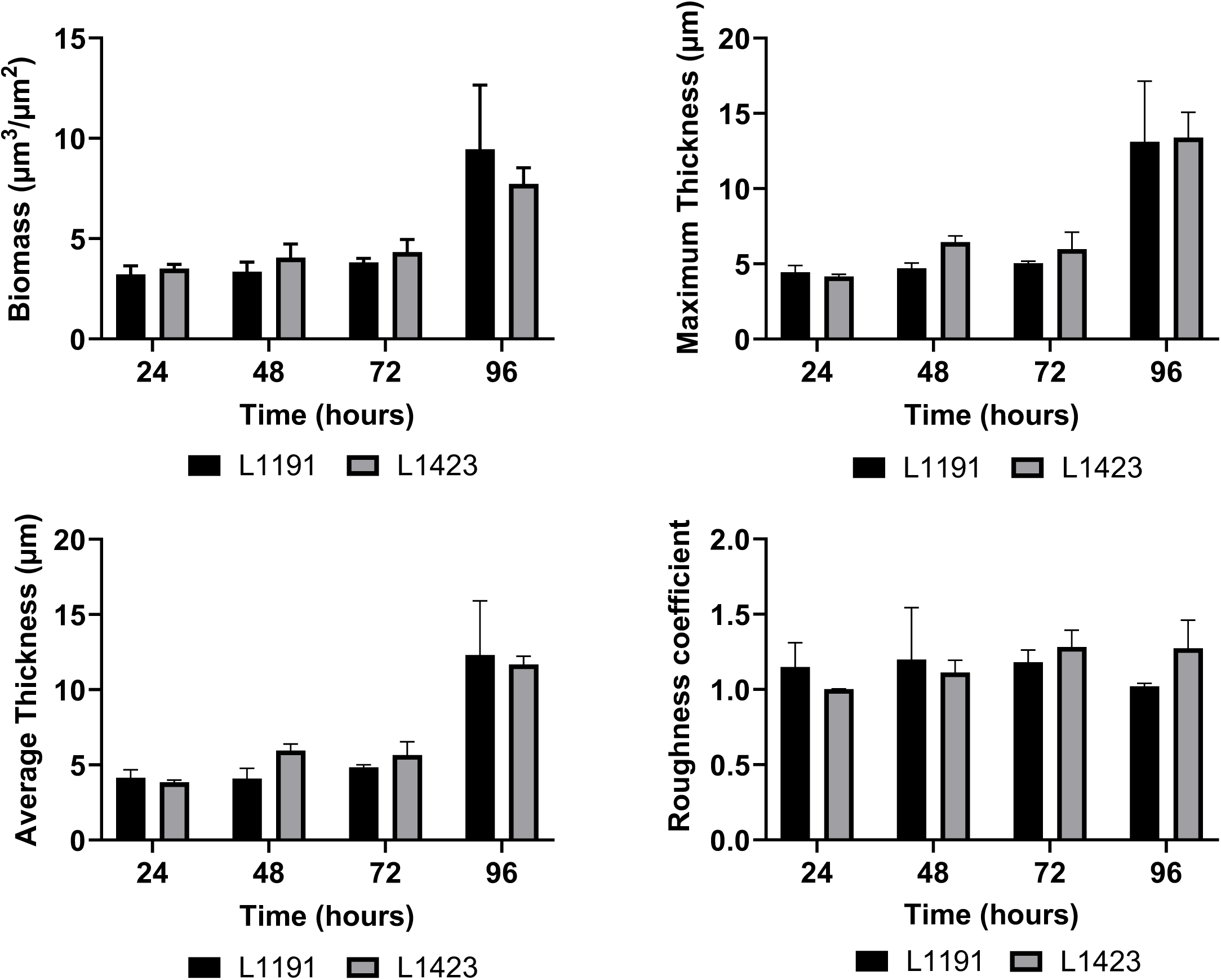
COMSTAT2 analysis of *B. pertussis* CLSM biofilm images. SNP cluster I strain, L1423, and SNP cluster II strain, L1191, were grown on glass cover slips and imaged at 24, 48, 72 and 96 h. Microscopy images were analysed using the COMSTAT2 ImageJ plugin. Biomass, maximum thickness and average thickness and roughness coefficient were calculated. The data are averages of three fields of view per biological replicate and three biological replicates were performed per strain per time point. Error bars represent standard deviation.

### Comparison of biofilm proteins between cluster I strain, L1423 and cluster II strain, L1191

To identify the underlying protein expression that may have led to the differences seen in the CLSM analysis, TMT-MS was performed on L1423 and L1191 biofilm extracted proteins. There were 1,453 proteins identified which is ∼45% of the total known *B. pertussis* proteins (Supplementary Table S2). A high proportion of proteins identified were predicted to be located in the cytoplasm (57%) while 14% of the proteins were located in the cytoplasmic membrane. The other proteins were distributed in the periplasmic space, outer membrane and extracellular space at 4%, 3% and 1%, respectively. The rest of the protein locations were unable to be predicted (19%). There were 40 proteins that were identified with significant (0.8 > FC > 1.2, adjusted *p* < 0.05) differences in expression between the two strains (Figure 3A). There were 18 proteins that were significantly downregulated and 22 significantly upregulated in L1423 compared to L1191 (Table 2 and Table 3).

**Figure 3.**
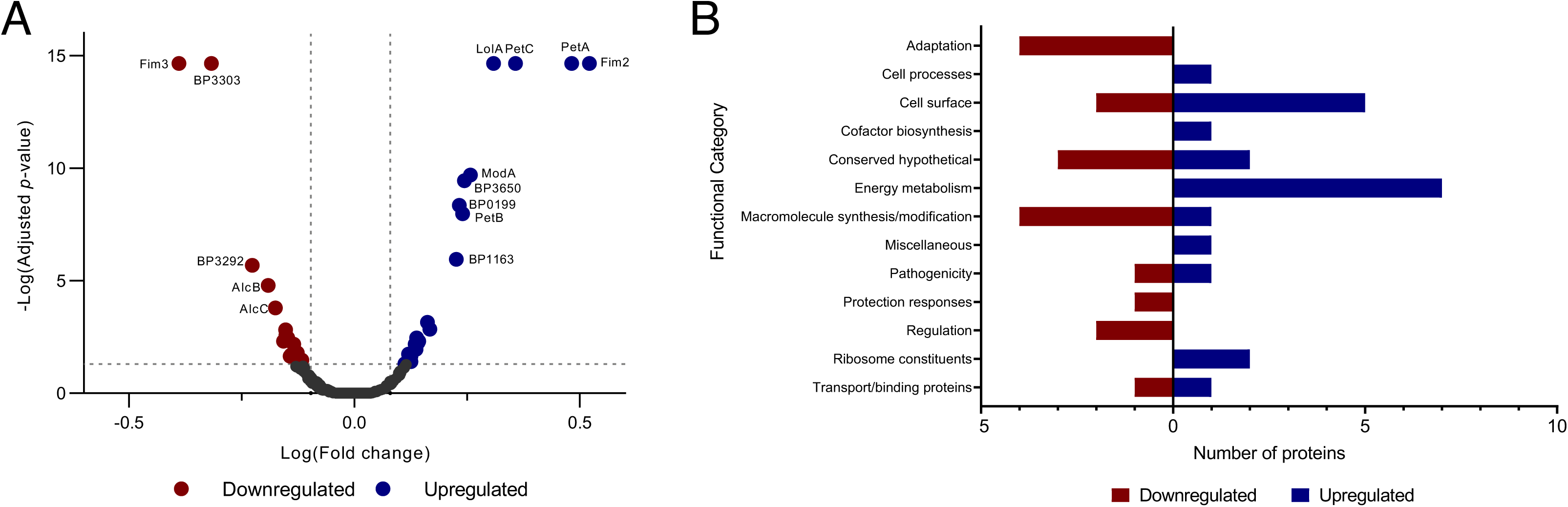
Proteins identified using TMT-MS between a representative *B. pertussis* SNP cluster I (L1423) and a SNP cluster II (L1191) strain. Proteins from a cluster I (L1423) and a cluster II (L1191) strain grown in biofilm conditions were extracted and measured by TMT-MS. **A** Volcano plot of proteins identified between the two strains. Proteins were deemed as upregulated with a fold change (FC) > 1.2, adjusted *p* < 0.05 and downregulated with a FC < 0.8, adjusted *p <* 0.05. Dashed grey lines mark 0.8 > FC > 1.2 and an adjusted *p* value = 0.05. Proteins with adjusted *p* < 0.001 are labelled on the plot. **B** Significantly differentially regulated proteins were grouped into functional categories based on Bart *et al*. (2). The number of proteins downregulated within each category are shown in red and the number of upregulated proteins are shown in blue.

**Table 1.**
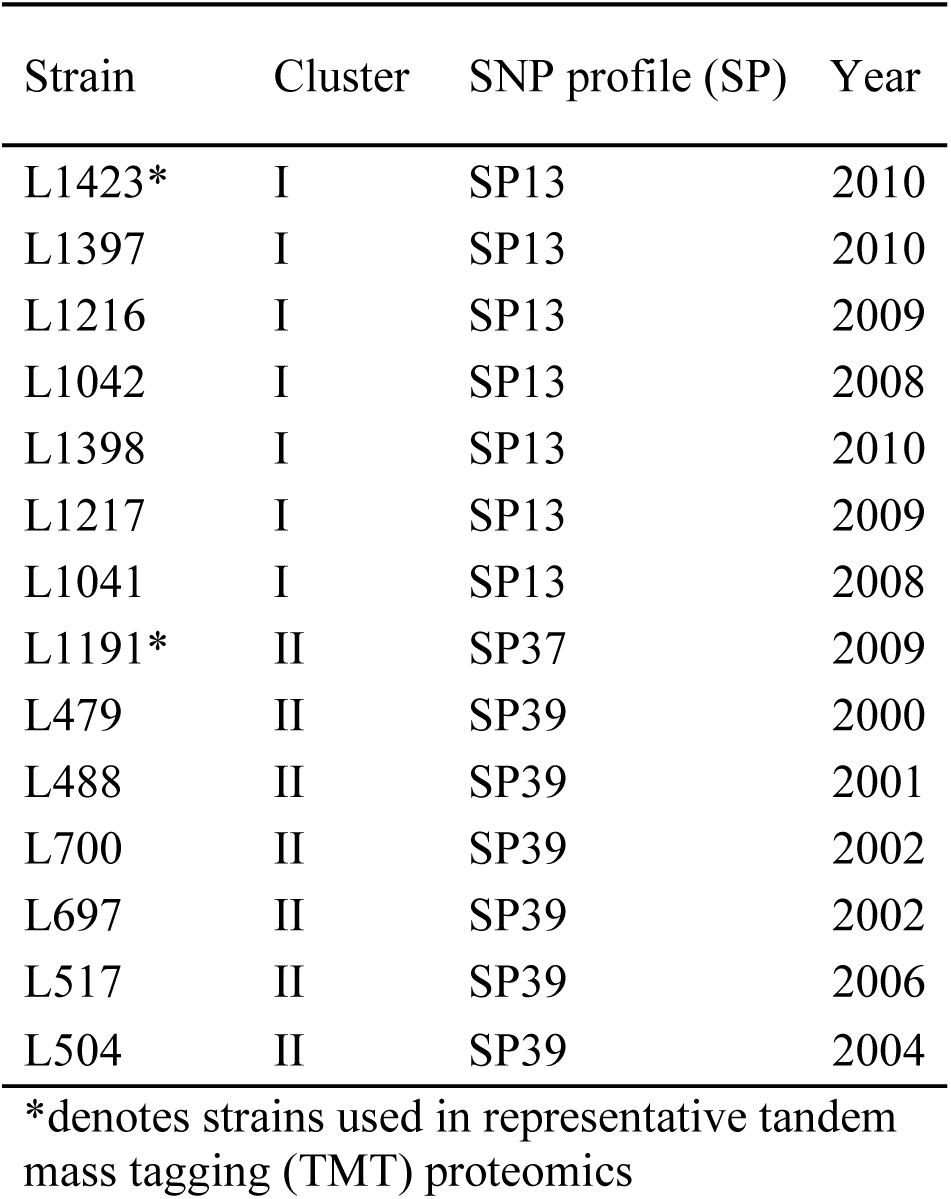
Strains used in this study.

**Table 2.**
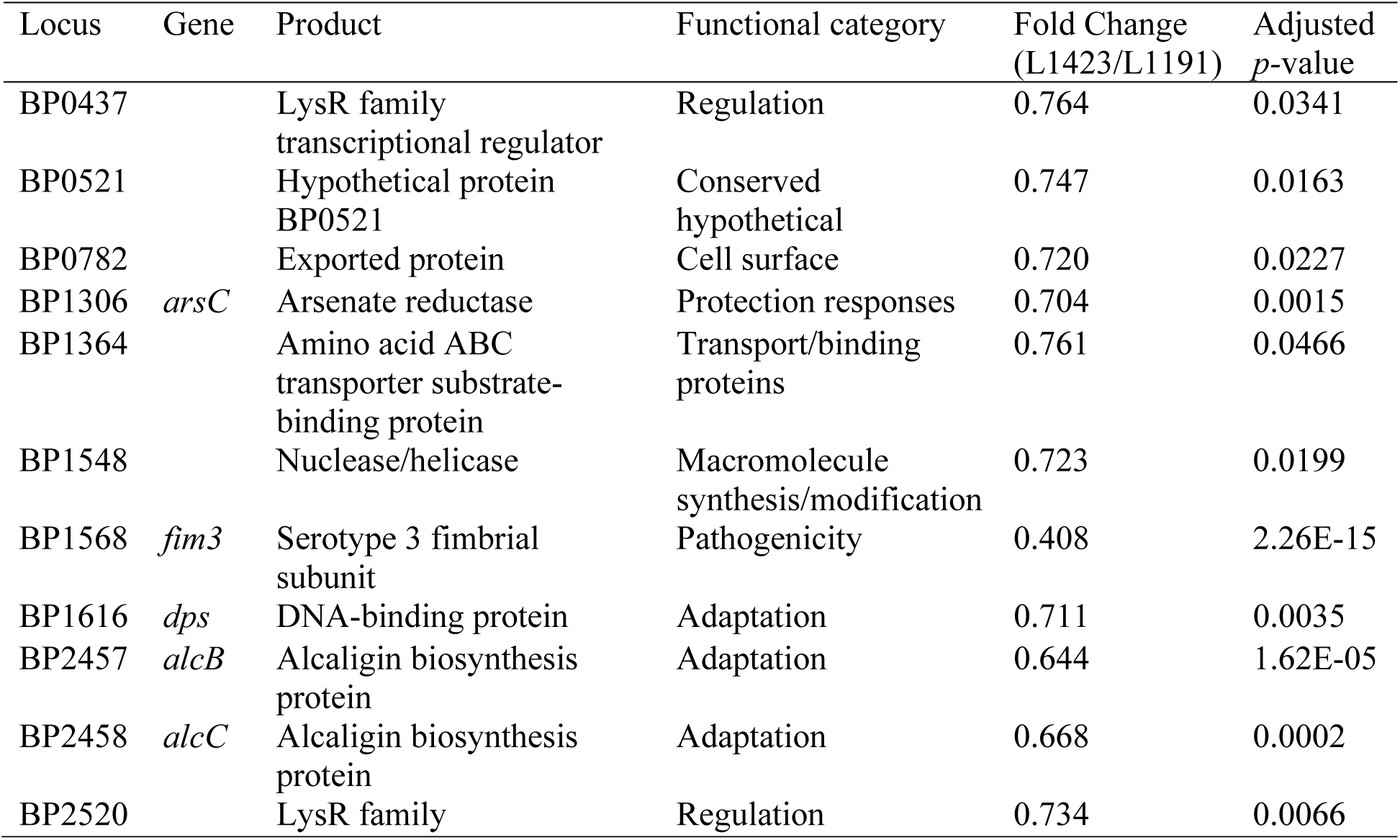
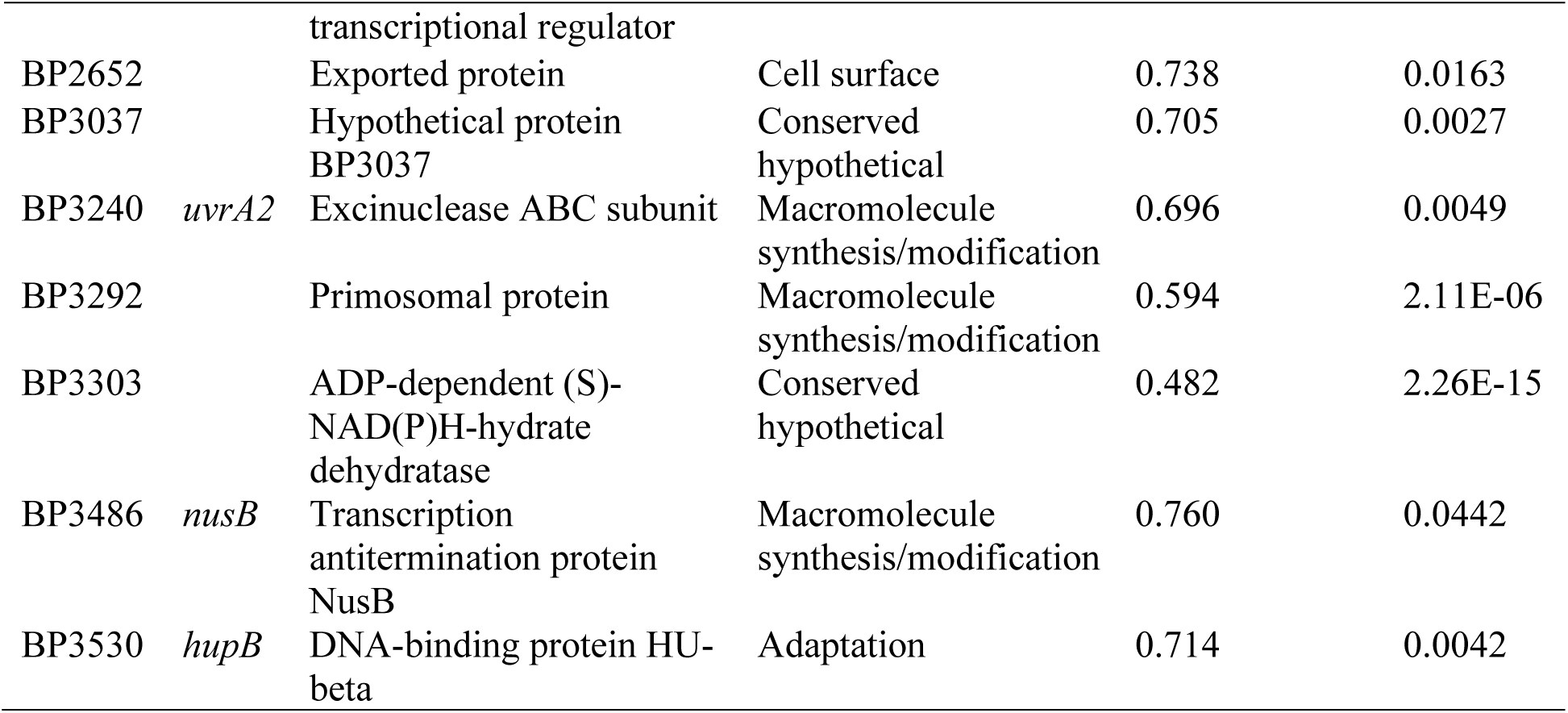
Proteins downregulated in L1423 *B. pertussis* biofilm cells compared to L1191.

**Table 3.**
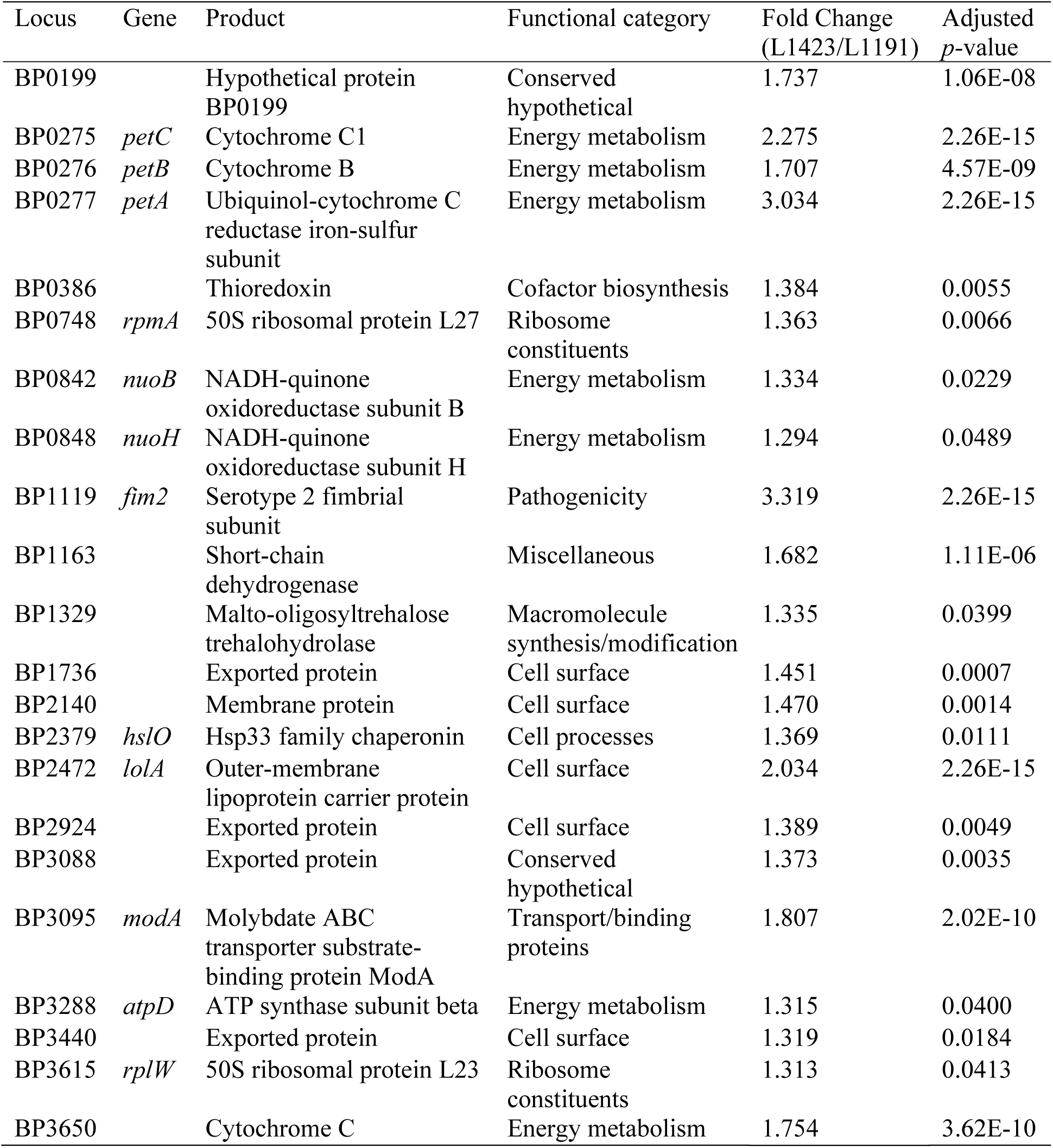
Proteins upregulated in L1423 *B. pertussis* biofilm cells compared to L1191.

The largest change was seen in proteins belonging to the energy metabolism functional category (Figure 3B). There were 7 proteins upregulated in energy metabolism in L1423. These include 4 cytochrome proteins (PetABC and BP3650) as well as NADH-quinone oxidoreductase subunits (NuoBH) and ATP synthase (AtpD). There were only two proteins related to pathogenicity that were differentially regulated: Fim2 upregulated in L1423 while Fim3 was upregulated in L1191 biofilm cells. Interestingly, two siderophore biosynthesis proteins, AlcBC, were downregulated in the L1423 biofilm cells. Proteins that were previously shown to have significant changes between the two strains in planktonic conditions (12), ModA and BP3303, were also seen to be both upregulated and downregulated in L1423, respectively. Two LysR transcriptional regulator proteins (BP0437 and BP2520) were downregulated as well as two DNA binding proteins (HupB and Dps).

Finally, there were 5 cell surface proteins that were upregulated in L1423 biofilm cells (Figure 3B).

### Confirmation of expression with MRM-hr

MRM-hr mass spectrometry was used to confirm the changes in expression identified in the TMT-MS experiment above. Of the 40 proteins with significantly differential expression between the strains, 9 proteins with suitable peptides were selected for confirmation with MRM-hr (Table 4 and Supplementary Table S1). Fim2 and Fim3 were excluded from the analysis as they were related to the strain serotype, which is already known (40). The upregulated proteins tested were: PetC, BP3650, BP2924, AtpD and RplW. The downregulated proteins tested were: AlcC, NusB, BP1364 and BP2652. Three non-differentially expressed proteins DnaK, ClpB and RpsA were selected as the control proteins for normalisation (Table 4).

**Table 4.** Proteins tested with MRM-hr for confirmation of TMT-MS results.

| Locus | Gene | Product | Functional Category | Fold change TMT | Adjusted *p*-value TMT |
| --- | --- | --- | --- | --- | --- |
| BP0275 | *petC* | Cytochrome C1 | Energy metabolism | 2.275 | 2.26E-15 |
| BP0950* | *rpsA* | 30S ribosomal protein S1 | Ribosome constituents | 0.961 | 0.997 |
| BP1198* | *clpB* | ATP-dependent protease, ATPase subunit | Macromolecule degradation | 0.945 | 0.997 |
| BP1364 |  | Amino acid ABC transporter substrate-binding protein | Transport/binding proteins | 0.761 | 0.047 |
| BP2458 | *alcC* | Alcaligin biosynthesis protein | Adaptation | 0.668 | 1.60E-04 |
| BP2499* | *dnaK* | Molecular chaperone DnaK | Cell processes | 0.924 | 0.997 |
| BP2652 |  | Exported protein | Cell surface | 0.738 | 0.016 |
| BP2924 |  | Exported protein | Cell surface | 1.389 | 0.005 |
| BP3288 | *atpD* | ATP synthase subunit beta | Energy metabolism | 1.315 | 0.040 |
| BP3486 | *nusB* | Transcription antitermination protein NusB | Macromolecule synthesis/modification | 0.760 | 0.044 |
| BP3615 | *rplW* | 50S ribosomal protein L23 | Ribosome constituents | 1.313 | 0.041 |
| BP3650 |  | Cytochrome C | Energy metabolism | 1.754 | 3.62E-10 |

\*denotes selection as control proteins for normalisation

The MRM-hr experiment confirmed PetC and BP3650 to be upregulated in L1423 while AlcC was confirmed to be downregulated in L1423 (adjusted *p*-value < 0.05) (Supplementary Table S3). AtpD was also seen to be downregulated in L1423 which was the opposite of the TMT result (Figure 4).

**Figure 4.**
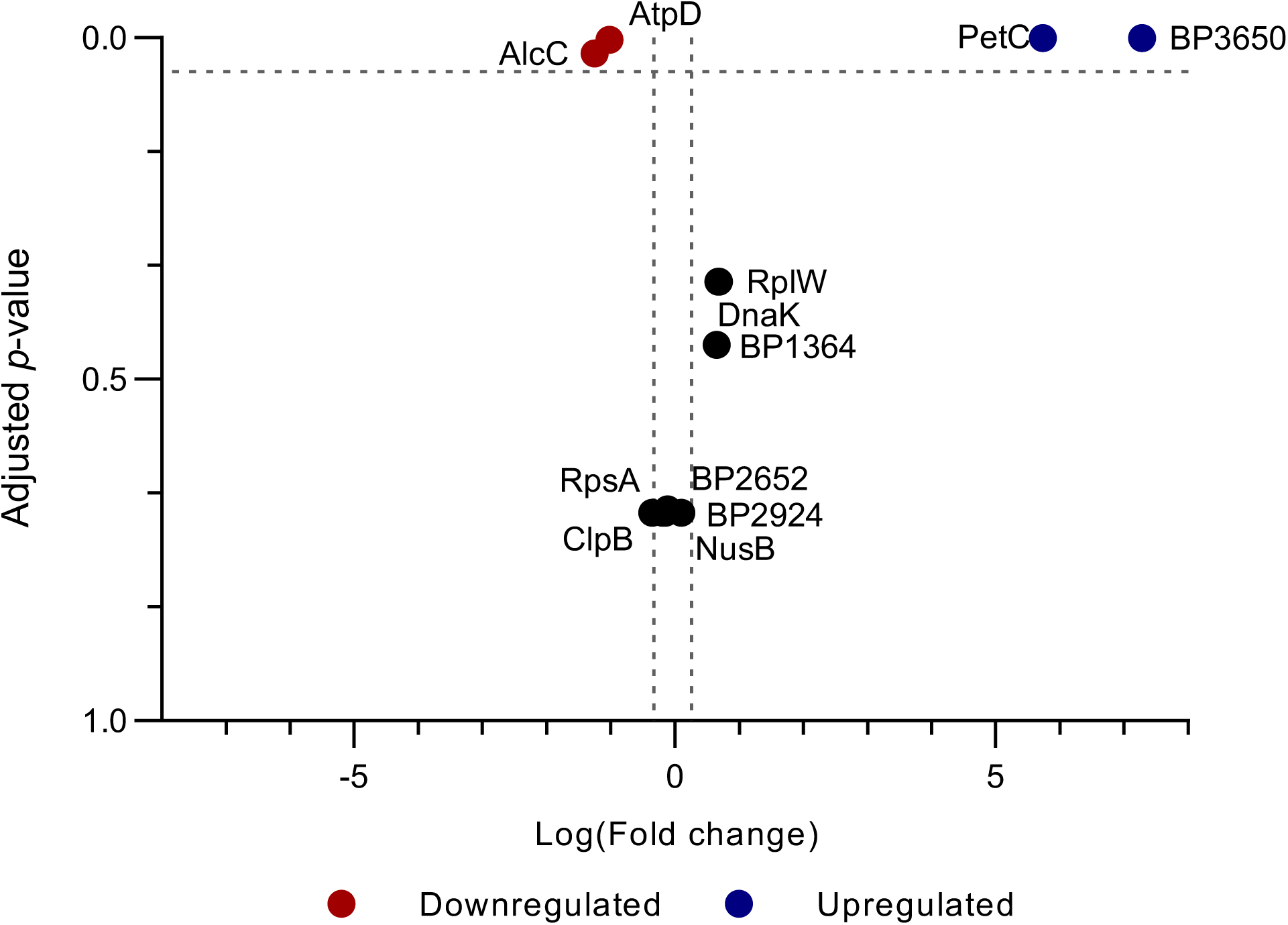
Confirmation of differentially expressed proteins identified in the TMT-MS using high resolution multiple reaction monitoring (MRM-hr). Volcano plot of nine proteins tested using MRM-hr. Three control proteins, DnaK, RpsA and ClpB, were chosen based on the TMT-MS results for normalisation. Proteins were deemed as upregulated with a fold change (FC) > 1.2, adjusted-*p* < 0.05 and downregulated with a FC < 0.8, adjusted *p <* 0.05. Dashed grey lines mark 0.8 < FC > 1.2 and an adjusted *p* value = 0.05.

Furthermore, to confirm whether the differences identified are strain differences or cluster differences, MRM-hr was performed on 6 additional strains from each cluster and the expression of the same 9 proteins mentioned above were measured. It was found that there were no significant differences in the expression of these proteins between the clusters (Supplementary Table S4). Interestingly, it was found that there was a high level of variability between strains (Figure 5). The proteins with the highest variation were ATP synthase (AtpD) and exported protein (BP2652) (CV = 27.08% and 27.04%) with L700 (SNP cluster II strain) and L1042 (SNP cluster II strain) with the highest abundance of these proteins, respectively. Notably, SNP cluster I strain L1398 had the lowest abundance for AtpD, BP2652 and an alcaligin biosynthesis protein (AlcC). However, L1398 had the highest value for a ribosomal protein (RplW) and relatively high values for cytochrome proteins, BP3650 and PetC (Figure 5).

**Figure 5.**
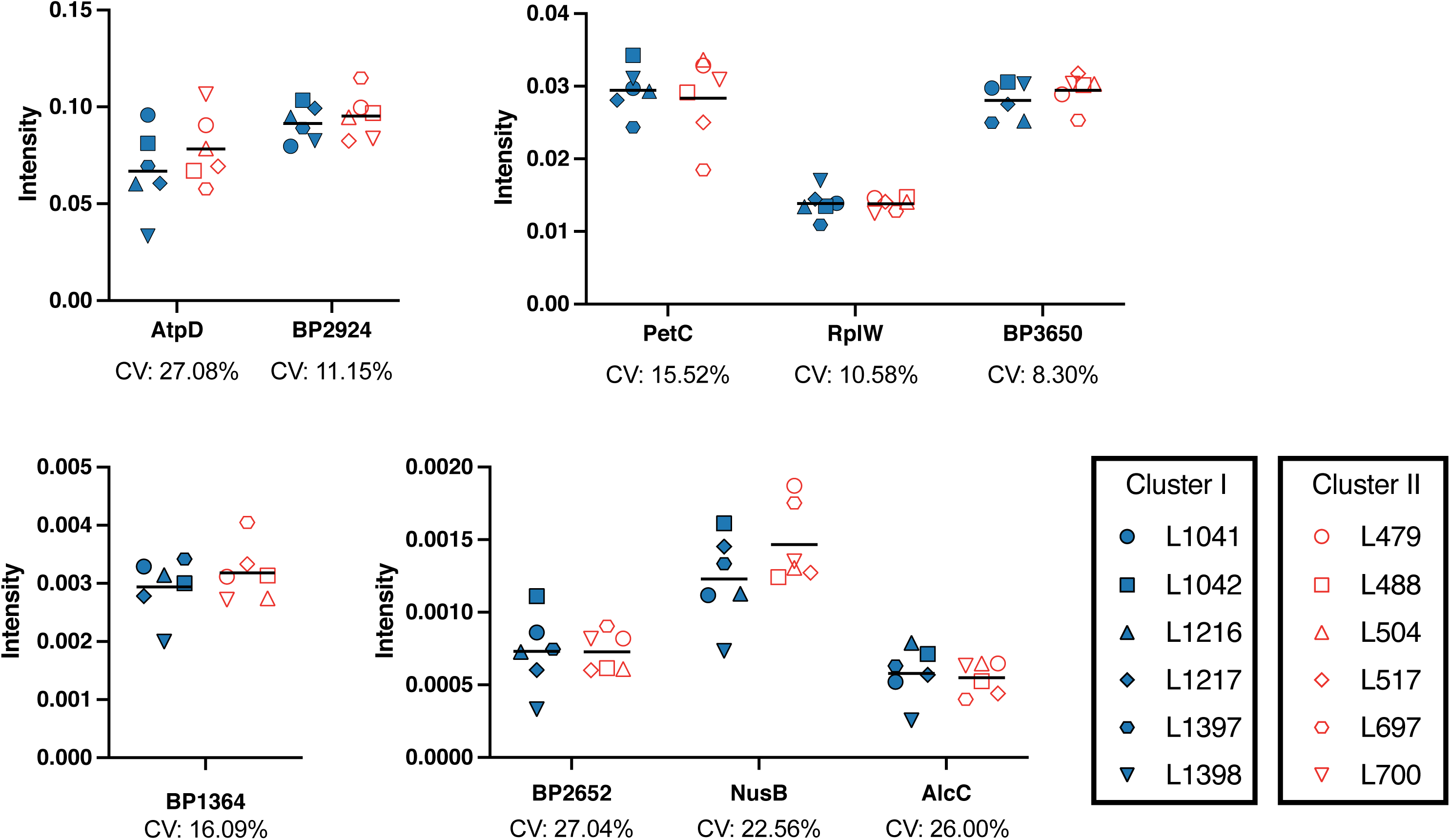
High resolution multiple reaction monitoring (MRM-hr) measurement of TMT-MS proteins of 6 additional strains from SNP cluster I and SNP cluster II. Relative abundances of proteins from individual strains were measured by MRM-hr. Six additional *B. pertussis* strains from both clusters were tested for a subset of the differentiated proteins identified from the representative strains. The clusters are represented as different colours and the strains as separate symbols. The protein intensity value (y-axis) is an average of all peptides measured for that protein for each strain. The bar indicates the average for the cluster. Coefficients of variation (CV) are indicated under the individual proteins. The proteins are grouped into separate graphs to allow differences in scale.

There were correlations in expression between particular proteins, suggesting that these proteins may be co-regulated. Expression of BP2652 was positively correlated with NusB (Pearson’s correlation: *r*(10) = 0.677, *p* = 0.016) and AtpD (*r*(10) = 0.606, *p* = 0.037) while negatively correlated with RplW (*r*(10) = -0.559, *p* = 0.039) (Figure 7). The expression of NusB was also positively correlated with BP2924 (*r*(10) = 0.726, *p* = 0.008). Finally, BP1364 was negatively correlated with RplW (*r*(10) = -0.638, *p* = 0.026) and PetC (*r*(10) = - 0.704, *p* = 0.011) (Figure 6).

**Figure 6.**
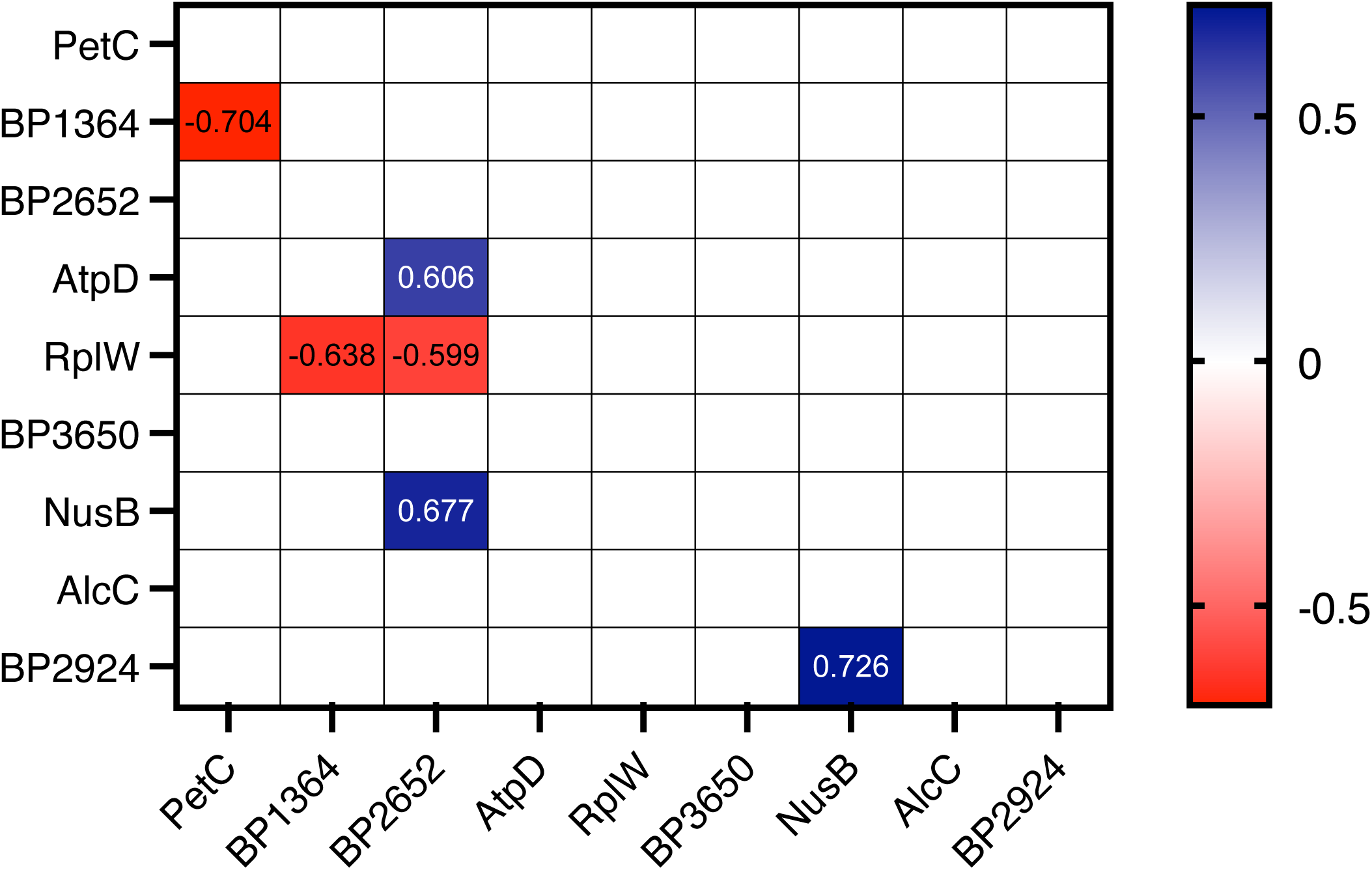
Pearson’s correlation matrix of significant correlations between protein expression measured with MRM-hr. Protein average intensity was measured from 12 individual strains of *B. pertussis* from both SNP cluster I and SNP cluster II. Protein average intensity is an average of all peptides measured for that protein. Significant changes are shown in the matrix. Colours relate to negative (red) and positive (blue) correlations. Proteins were considered positively correlated if *r* > 0.5, *p* < 0.05 and negatively correlated if *r* < -0.5, *p* < 0.05.

## Discussion

Despite high vaccination coverage, there has been a resurgence in *B. pertussis* notifications globally (3, 41–44). It has been hypothesised that a contributing factor to the resurgence is related to the ability of *B. pertussis* to form biofilms (16, 18). Increased biofilm formation capability has previously been hypothesised to increase respiratory tract colonisation in clinical isolates (20). Also related to the resurgence is the expansion of SNP cluster I strains (*ptxP3* allele type) dominating over the previously dominant SNP cluster II strains (*ptxP1* allele type) (3, 24, 45, 46). Previous studies have examined differences between strains from these clusters in planktonic conditions (11, 12, 23). In this study, we compared cluster I and cluster II strains in biofilm formation and protein expression to identify changes that may help explain the cluster I dominance. We found that the two SNP clusters had similar biofilm formation capabilities, but their biofilm structures were different.

Two representative strains from the clusters had consistent growth of biofilm and comparable levels of biomass at 96 h (Figure 2). However, the cluster II strain was observed to have a more diffuse structure while the cluster I strain had a denser biofilm structure (Figure 1). Differences in *B. pertussis* mature biofilm structure have been reported between strains from two different regions (Argentina and USA) (20). It was found that the biofilms from strains isolated from Argentina made uniform layers of cells similar to those identified for both strains in this study while the strains from USA formed irregularly shaped aggregates (20). It should be noted that the structural differences identified between the Argentinian and USA *B. pertussis* strains (20) are more easily recognisable compared to those identified in this study. While these studies identified differences on the substratum level of bound biofilm (i.e., biofilms covering the entire surface vs clustered aggregates) the changes identified in this study relate to the density of the cells within the biofilm. The results presented in this study suggest that if the dominance of cluster I strains were related to biofilm, there are more subtle changes rather than overall biofilm formation capabilities that may contribute to the increased fitness previously observed (9).

This study explored differences in protein expression between cluster I and cluster II strains in biofilm conditions to identify expression changes that may be related to the differences in structure. The most significant change between the two representative strains were upregulation of PetABC and BP3650 cytochrome proteins. PetABC is the cytochrome bc complex (complex III) that transfers electrons to cytochrome c (BP3650) and translocates protons across the membrane to create the proton motive force that powers ATP synthase (47). PetABC was found to be upregulated in iron repressed conditions and in low oxygen conditions in other species (48–50). This suggest that the regulatory pathways that are induced in low iron or oxygen may be triggered in the cluster I strain. A low iron and oxygen environment could be related to a lower diffusion of the nutrients in the cluster I strain due to the denser biofilm structure identified in the confocal images.

Analysis of additional strains found that PetABC and BP3650 changes were strain specific to L1191 (Figure 5). This may be explained by a genomic change that was observed specifically in the cluster II L1191 strain. The sequences of L423 and L1191 were aligned, and an insertion sequence (IS*481*) was found in the *petB* gene (nucleotide position 619920) in L1191 (Supplementary Figure S1). This insertion may disrupt the protein so that it is not produced correctly. The protein sequences were aligned to observe changes that may occur in the protein function. It was found that the IS led to a truncation (amino acid position 418) of the C-terminus which includes a predicted transmembrane domain of the PetB protein which may affect its function as an electron transport chain complex (Supplementary Figure S2). This disruption may also lead to the changes observed in BP3650 expression as they are both linked as cytochrome proteins. While not a cluster specific change, it is noteworthy that this change in expression was not observed when the strains were compared planktonically (12). This suggests that the PetABC and BP3650 proteins may have a specific regulator that is only triggered in biofilm conditions for *B. pertussis*.

Our previous study had compared the same strains used in this study in planktonic conditions (12). Any overlap between the two studies would likely identify changes that are consistent between the two clusters regardless of the condition (biofilm or planktonic). There were 5 proteins identified that had the same protein expression trends between the studies. Of these, the downregulation of BP3303 and upregulation of ModA in the cluster I strain may be as a result of genomic changes between the strains. Both genes encoding these proteins have non-synonymous SNPs as mentioned in Luu *et al.* (12). BP3303 contains a SNP in L1423 at nucleotide position 699 causing a change from glycine to arginine. This SNP has been found in 19/22 of the cluster I strains sequenced from the Australian 2008 – 2012 epidemic (40). It is a hypothetical protein which requires further studies to elucidate its role. Adjacent to *modA*, *modB* contains a SNP in L1191 at nucleotide position 29 causing a change from isoleucine to threonine. ModABC has been demonstrated to have a role in chronic lung infection and biofilm formation in *P. aeruginosa* (51, 52). The SNPs have been predicted to lead to reduced protein stability, however the effect on protein functionality have not been experimentally determined (12). *modB* has also been shown to have increased expression in *ptxP3* strains compared to *ptxP1* in sulfate modulated conditions in a previous study (24).

These studies highlight a potential link between the expression of ModABC and the dominance of the cluster I strains. There were 3 additional proteins, Fim2, Fim3 and BP2924, identified in this study that reflected the results found in the planktonic comparison (12). Fim2 and BP2924 were upregulated while Fim3 was downregulated in the cluster I strain. The cluster I strain is a Fim2 serotype while the cluster II strain is a Fim3 serotype, therefore, these results confirm the expected changes in protein expression (40). BP2924 is a hypothetical cell surface protein that was also found to be upregulated in the secretome of the same cluster I strain used in this study although it was unable to be confirmed using MRM-hr (35). Furthermore, *BP2924* was identified as transcriptionally upregulated in the Dutch *ptxP3* strains compared to the *ptxP1* strains (4). Therefore, BP2924 may be linked to a competitive advantage for the cluster I strains.

It has been well established that the current SNP cluster I (*ptxP3*) strains have an advantage over the previously dominant SNP cluster II (*ptxP1*) strains (4, 9, 10, 12). Due to the low genetic diversity in *B. pertussis* evolution, it is expected that additional strains from the cluster would reflect the changes identified in the representative strains (7, 8). Many of the changes identified in this study were unable to be confirmed using the MRM-hr method in the additional cluster strains due to lack of suitable peptides. Of the 40 proteins identified as differentially regulated between the representative strains, only 9 could be examined and were shown to have variable expression for the additional strains included from each cluster (Figure 5). Interestingly, it was seen that a collection of proteins displayed correlated trends in expression between strains regardless of clusters (Figure 6). This suggests that these proteins may be co-expressed or that the expression of one affects the other. Nevertheless, these results suggests that the differences are likely strain specific differences rather than cluster specific differences. Previous studies have shown that the variability in gene expression between previously circulating and currently circulating strains is very small when considering the total number of coding genes (4, 12, 13, 24). With the exclusion of Fim2 and Fim3, there was an additional 29 proteins that were identified in the representative strains with differential expression that were to be confirmed. These additional 29 proteins may still be cluster specific changes and would require further investigation. Included within these proteins was a downregulation of two LysR transcriptional regulators and two DNA binding proteins, HupB and Dps, in the cluster I strain. There is little research in the regulatory role of these proteins in *B. pertussis* and studies have shown that there are regulators that control the expression of a wide range of *B. pertussis* genes and proteins (53–58). It is possible that these regulators are linked to the difference in biofilm structure and behaviour between L1423 and L1191.

In conclusion, this study found that there were no differences in the amount of biofilm biomass produced between cluster I and cluster II strains, suggesting that biofilm formation capability may not be a contributing factor to the fitness of the currently circulating cluster I strains over cluster II strains. However, there were differences in mature biofilm structure between the representative strains, which warrants further investigation. Understanding the underlying protein expression that triggers these differences in biofilm behaviour may provide insight into the biofilm formation process of *B. pertussis*. Differential expression of BP3303, ModA and BP2924 found in other studies (4, 24, 59) were confirmed to have the same trend in biofilm conditions between the cluster I and cluster II strains. The replicability of these findings suggests a potentially important role of these proteins in the dominance of cluster I strains. Finally, this study highlights that proteomic changes in biofilm conditions between clusters are compounded by strain specific differences, underscoring the proteomic diversity of the *B. pertussis* population and contrasting the genetic homogeneity of the species (7, 8).

## Acknowledgements

Hiroki Suyama was supported by the Australian Government Research Training Program Scholarship. This work was supported by a grant from the National Health and Medical Research Council of Australia.

## Data availability

The mass spectrometry proteomics data have been deposited to the ProteomeXchange Consortium via the PRIDE (51) partner repository with the dataset identifier PXD046513.

